## Supplemental figures for "Ablation of STAT3 in Purkinje Cells Reorganizes Cerebellar Synaptic Plasticity in Long-Term Fear Memory Network"

**Table S1. List of up/down genes from analysis of KEGG pathway**

| **log(pval)** | **KEGG pathway, upgenes** |  | **log(pval)** | **KEGG pathway, down genes** |
| --- | --- | --- | --- | --- |
| 22.26281 | Alzheimer's disease |  | 4.017729 | Phototransduction |
| 18.08619 | Parkinson's disease |  | 2.165579 | Neuroactive ligand-receptor interaction |
| 18.03716 | Oxidative phosphorylation |  |  |  |
| 14.66555 | Huntington's disease |  |  |  |
| 10.31785 | Non-alcoholic fatty liver disease (NAFLD) |  |  |  |
| 10.14388 | Retrograde endocannabinoid signaling |  |  |  |
| 8.869666 | Synaptic vesicle cycle |  |  |  |
| 8.580044 | Dopaminergic synapse |  |  |  |
| 8.291579 | Ribosome |  |  |  |
| 7.019088 | Adrenergic signaling in cardiomyocytes |  |  |  |
| 6.962574 | Cardiac muscle contraction |  |  |  |
| 6.419075 | Long-term depression |  |  |  |
| 5.524329 | GABAergic synapse |  |  |  |
| 5.374688 | Long-term potentiation |  |  |  |
| 5.179799 | Glutamatergic synapse |  |  |  |
| 5.175224 | Citrate cycle (TCA cycle) |  |  |  |
| 5.084073 | Endocrine and other factor-regulated calcium reabsorption |  |  |  |
| 4.899629 | Thyroid hormone signaling pathway |  |  |  |
| 4.752027 | cGMP-PKG signaling pathway |  |  |  |
| 4.716699 | Carbon metabolism |  |  |  |
| 4.182435 | Endocytosis |  |  |  |
| 3.742321 | Oxytocin signaling pathway |  |  |  |
| 3.514279 | Sphingolipid signaling pathway |  |  |  |
| 3.293282 | Amphetamine addiction |  |  |  |
| 3.291579 | Ubiquitin mediated proteolysis |  |  |  |
| 3.273273 | cAMP signaling pathway |  |  |  |
| 3.109579 | Autophagy - animal |  |  |  |
| 3.094744 | Phosphatidylinositol signaling system |  |  |  |
| 3.067526 | Biosynthesis of amino acids |  |  |  |
| 3.013676 | Glucagon signaling pathway |  |  |  |
| 2.970616 | Salivary secretion |  |  |  |
| 2.847712 | Oocyte meiosis |  |  |  |
| 2.707744 | Insulin secretion |  |  |  |
| 2.694649 | Gastric acid secretion |  |  |  |
| 2.517126 | Choline metabolism in cancer |  |  |  |
| 2.333482 | Circadian entrainment |  |  |  |
| 2.117475 | Fc gamma R-mediated phagocytosis |  |  |  |
| 2.091515 | Metabolic pathways |  |  |  |
| 2.06956 | Ferroptosis |  |  |  |
| 1.931814 | Estrogen signaling pathway |  |  |  |
| 1.876148 | Collecting duct acid secretion |  |  |  |
| 1.707744 | Gap junction |  |  |  |
| 1.642065 | 2-Oxocarboxylic acid metabolism |  |  |  |
| 1.595166 | Mitophagy - animal |  |  |  |
| 1.531653 | Pancreatic secretion |  |  |  |
| 1.496209 | Calcium signaling pathway |  |  |  |
| 1.413413 | Salmonella infection |  |  |  |
| 1.376751 | Inositol phosphate metabolism |  |  |  |
| 1.374688 | Focal adhesion |  |  |  |

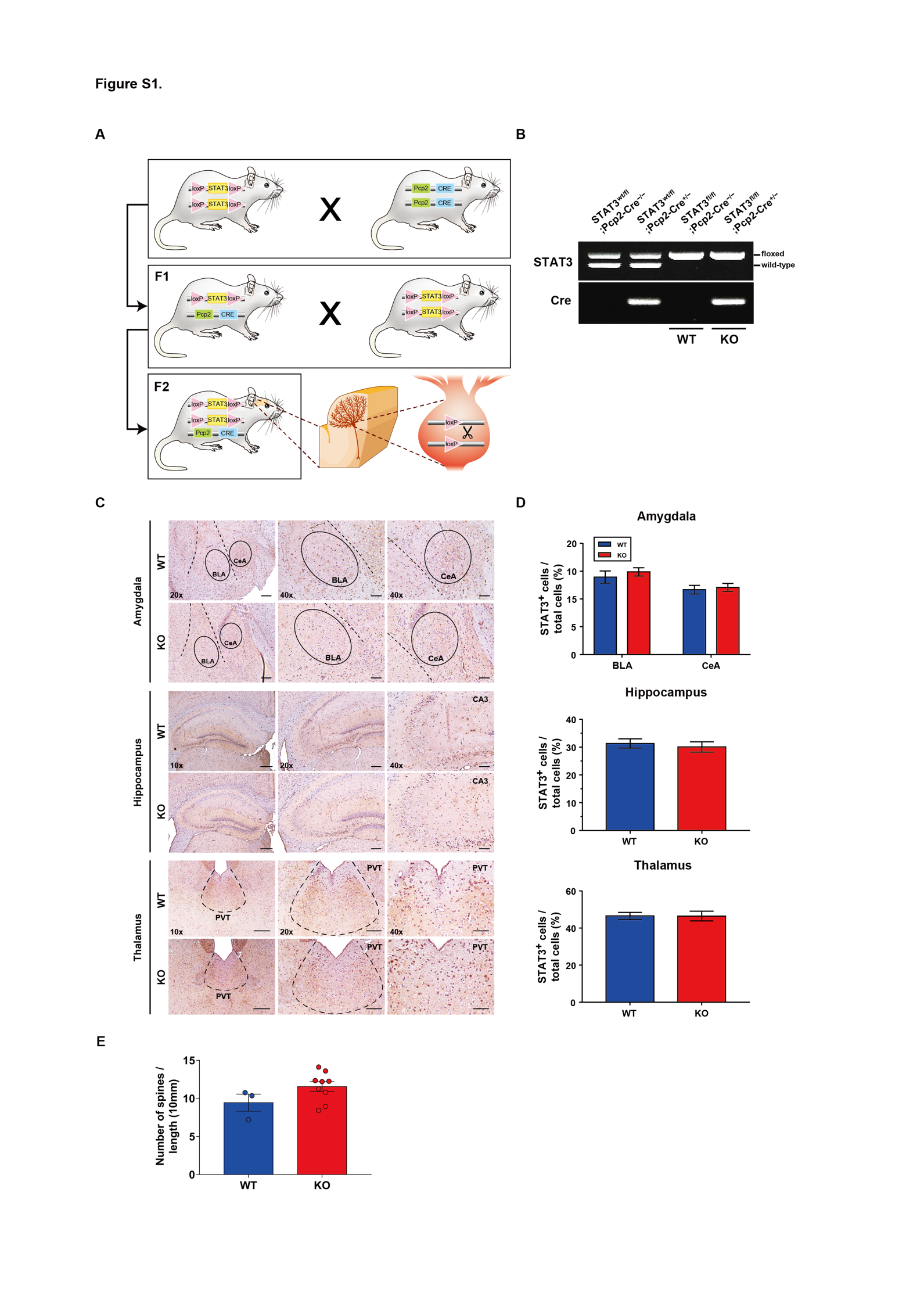

**Figure S1. Generation of STAT3^PKO^ mice model**

(A) Schematic representation of a breeding strategy for Purkinje cell-specific STAT3 knockout mouse generation. The first-generation offspring (F1) were obtained from STAT3 floxed mice crossing Pcp2-Cre mice, and STAT3^wt/fl^;Pcp2-Cre^−/−^ (F1) were further mated with STAT3 floxed mice. (B) Genotype analysis of the second-generation offspring (F2) using RT-PCR. The second-generation offspring (F2) were divided into four genotypes: STAT3^wt/fl^;Pcp2-Cre^−/−^, STAT3^wt/fl^;Pcp2-Cre^+/−^, STAT3^fl/fl^;Pcp2-Cre^−/−^ (WT), STAT3^fl/fl^;Pcp2-Cre^+/−^ (KO). (C) Immunohistochemistry analysis for STAT3 in amygdala, hippocampus, and thalamus regions from WT and STAT3^PKO^ mice. Scale bars of 10X image = 200 μm, 20X image = 100 μm, and 40X image = 50 μm. (D) Bar graphs show quantification of STAT3 expression (WT vs. STAT3^PKO^, Amygdala-BLA: 13.9 ± 1.09 vs. 14.8 ± 0.737, *p*=0.487; Amygdala-CeA: 11.6 ± 0.773 vs. 12.0 ± 0.705, *p*=0.702; Hippocampus: 31.3 ± 1.66 vs. 30.0 ± 1.85, *p*=0.624; Thalamus: 46.5 ± 1.91 vs. 46.4 ± 2.61, *p*=0.985, n = 8 slices of 5 mice per experimental group; two-tailed Student’s *t* test). (E) Measurement for the number of dendritic spines in Purkinje cells WT vs. STAT3^PKO^, 9.44 ± 1.12 vs. 11.5 ± 0.640, *p*=0.131, n =3, 9 cells; two-tailed Student’s *t* test). Data are presented as mean ± SEM.

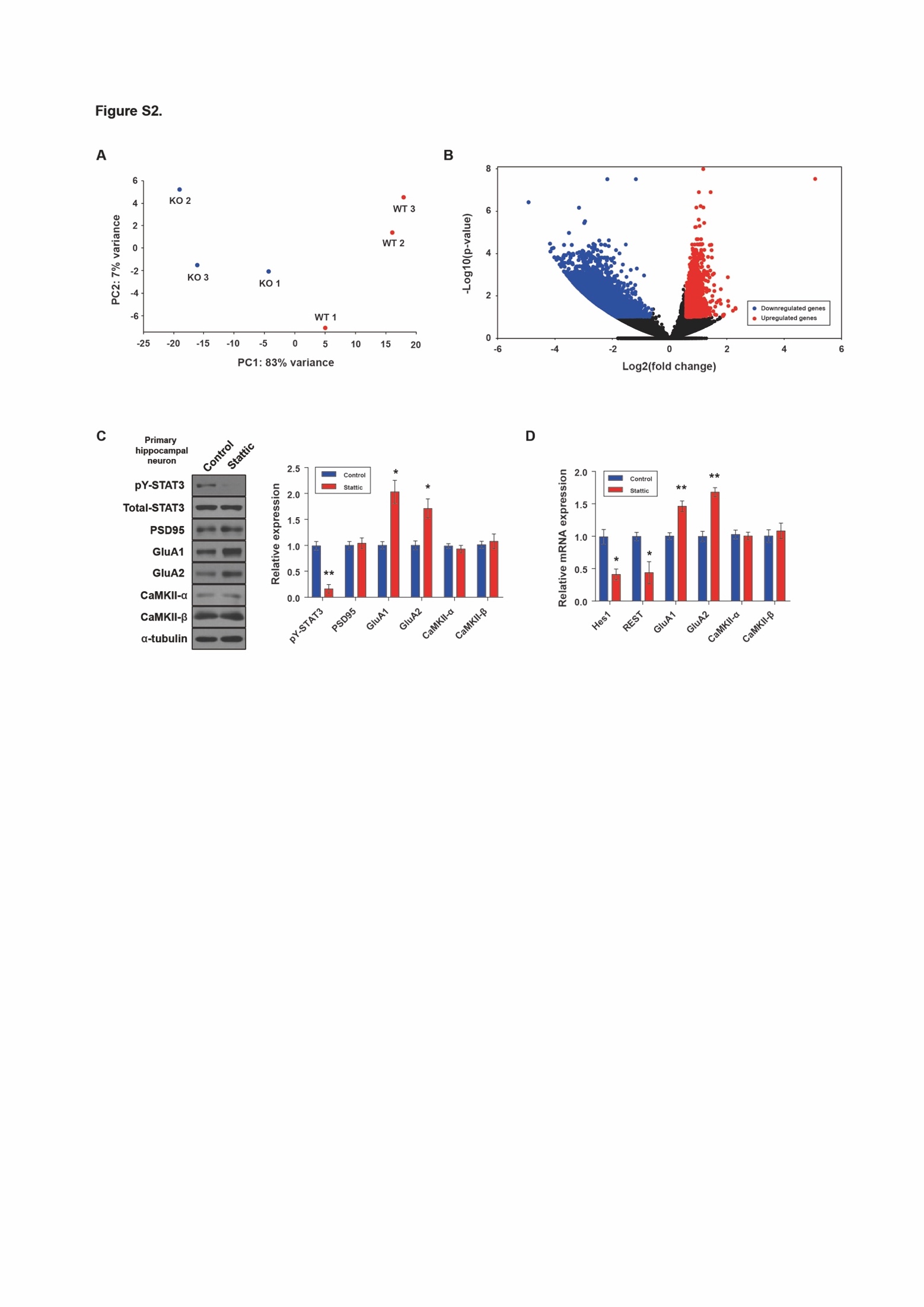

**Figure S2. Transcriptome analysis of Purkinje cells in WT and STAT3^PKO^ mice, and mechanism underlying the STAT3-modulated AMPA receptor expressions in hippocampal neuron model.**

(A) Principal component analysis of the normalized RNA-seq data for WT and STAT^PKO^ groups. (B) Volcano plot representation of fold change of transcripts in WT and STAT^PKO^ groups, *p*<0.05. (C) Western blot analysis of pY-STAT3, STAT3, PSD95, GluA1, GluA2, CaMKII-α, CaMKII-β, and α-tubulin in primary hippocampal neuron with or without STAT3 inhibition (STAT3: *p*=0.00220, GluA1: *p*=0.0120, GluA2: *p*=0.0254, n=3 mice; two-tailed Student’s *t* test). Quantification of Western blot analysis was obtained with relative densitometry and normalized with α-tubulin. (D) Relative mRNA levels of Hes1, REST, GluA1, GluA2, CaMKII-α, and CaMKII-β in primary hippocampal neuron with or without STAT3 inhibition (Hes1: *p*=0.0149, REST: *p*=0.0364, GluA1: *p*=0.00870, GluA2: *p*=0.00300, n=3 mice; two-tailed Student’s *t* test). Data are presented as mean ± SEM, and ^*^*p*<0.05, ^**^*p*<0.01.

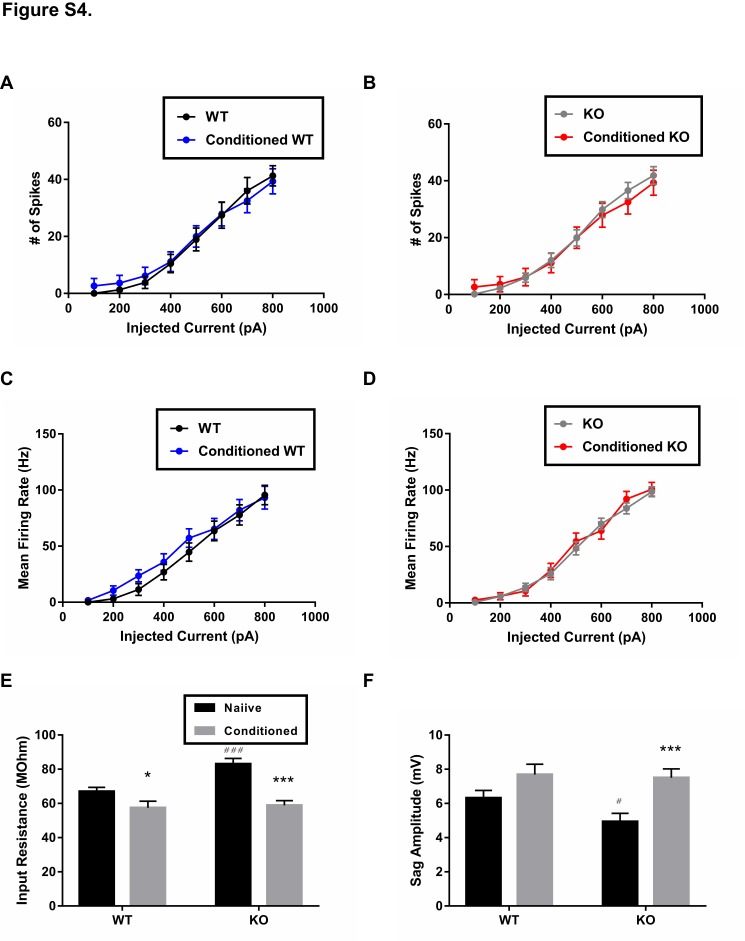

**Figure S3. Whole cell current-clamp recordings for measuring intrinsic excitability of PC in WT and STAT3^PKO^ mice. (n=24, 22, 29, 22 cells; WT, Conditioned WT, KO, and Conditioned KO groups, respectively).**

(A) Measurement for number of spikes when PCs in WT mice were injected a series of increasing current steps at 100 pA intervals, before and after fear conditioning (genotype × injected current interaction: F_(7,352)_=0.0655, *p*=0.999; genotype effect: F_(1,352)_=0.688, *p*=0.407; injected current effect: F_(7,352)_=48.1, *p*<0.0001; WT and conditioned WT groups, respectively; two-way ANOVA with Bonferroni correction). (B) Measurement for number of spikes when PCs in KO mice were injected a series of increasing current steps at 100 pA intervals, before and after fear conditioning (genotype × injected current interaction: F_(7,392)_=0.129, *p*=0.996; genotype effect: F_(1,392)_=3.50, *p*=0.0619; injected current effect: F_(7,392)_=69.1, *p*<0.0001; KO and conditioned KO groups, respectively; two-way ANOVA with Bonferroni correction). (C) Measurement for mean firing rate when PCs in WT mice were injected a series of increasing current steps at 100 pA intervals, before and after fear conditioning (genotype × injected current interaction: F_(7,360)_=0.2741, *p*=0.963; genotype effect: F_(1,360)_=2.55, *p*=0.110; injected current effect: F_(7,360)_=45.7, *p*<0.0001; WT and conditioned WT groups, respectively; two-way ANOVA with Bonferroni correction). (D) Measurement for mean firing rate when PCs in KO mice were injected a series of increasing current steps at 100 pA intervals, before and after fear conditioning (genotype × injected current interaction: F_(7,400)_=0.504, *p*=0.831; genotype effect: F_(1,400)_=0.0226, *p*=0.880; injected current effect: F_(7,400)_=125, *p*<0.0001; KO and conditioned KO groups, respectively; two-way ANOVA with Bonferroni correction). (E) Measurement for input resistance at -500 pA in WT and STAT^PKO^ mice, before and after fear conditioning (*p*<0.05, WT vs. Conditioned WT, two-tailed Student’s *t* test; *p*<0.001, WT vs. KO, two-tailed Student’s *t* test; *p*<0.001, KO vs. Conditioned KO, two-tailed Student’s *t* test). (F) Measurement for Sag amplitude in WT and STAT^PKO^ mice, before and after fear conditioning (*p*=0.0735, WT vs. Conditioned WT, two-tailed Student’s *t* test; *p*<0.05, WT vs. KO, two-tailed Student’s *t* test; *p*<0.001, KO vs. Conditioned KO, two-tailed Student’s *t* test). Data are presented as mean ± SEM, and ^*^*p*<0.05, ^**^*p*<0.01, ^***^*p*<0.001, compared with naïve group; ^#^*p*<0.05, ^###^*p*<0.001, compared with naïve WT group.

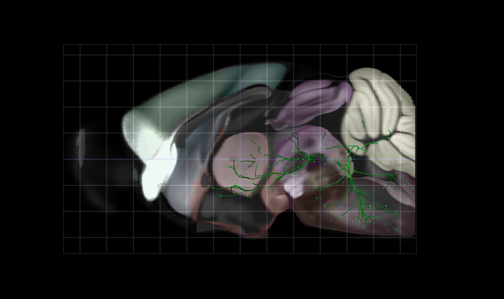

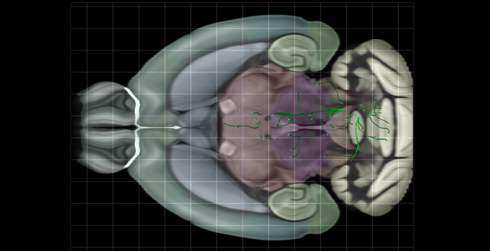

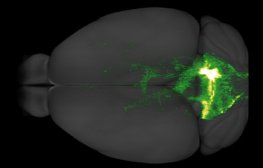

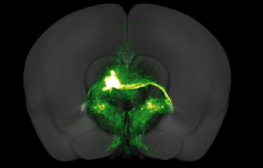

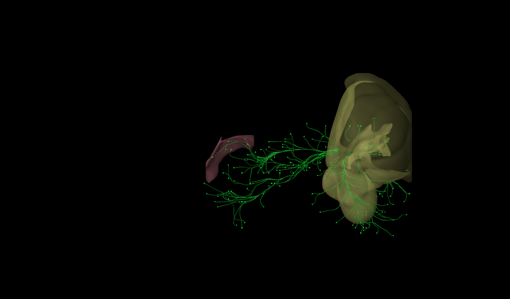

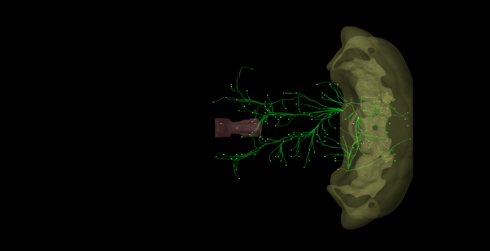

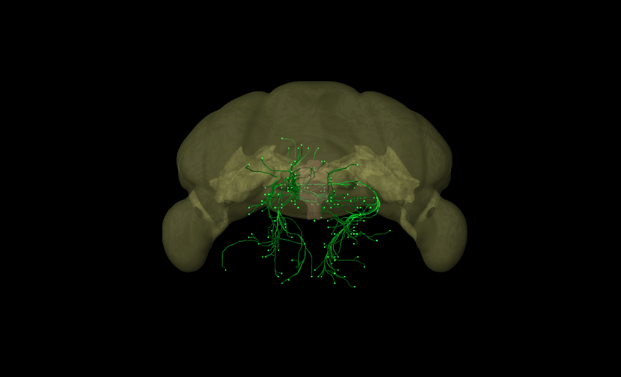

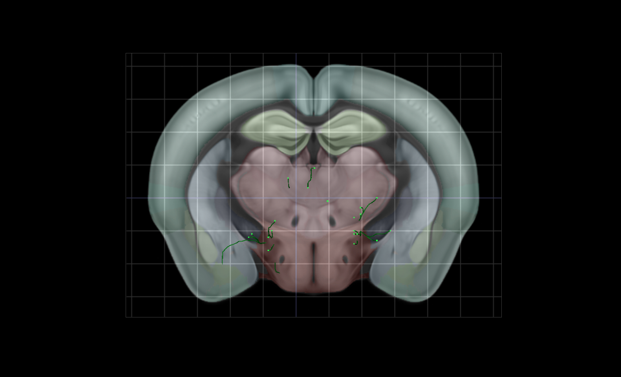

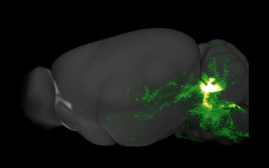

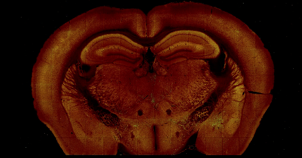

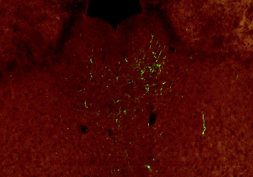

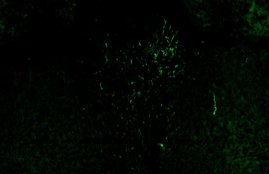

**A.**

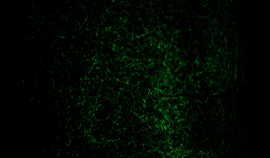

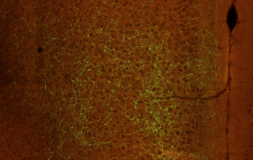

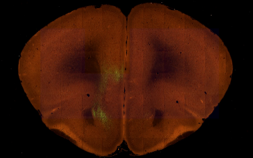

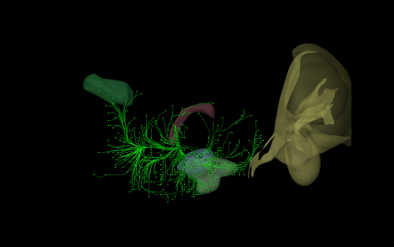

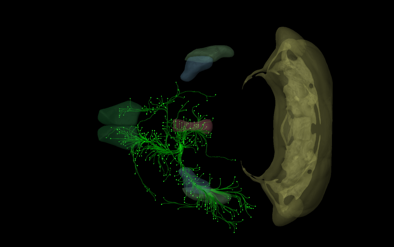

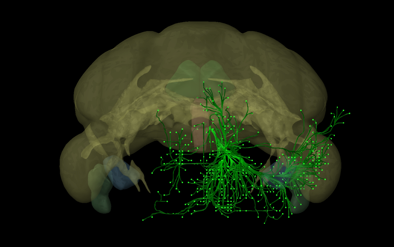

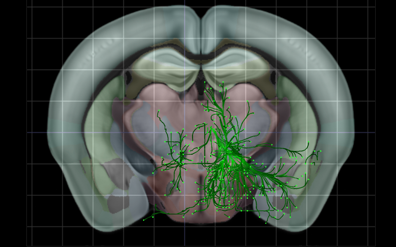

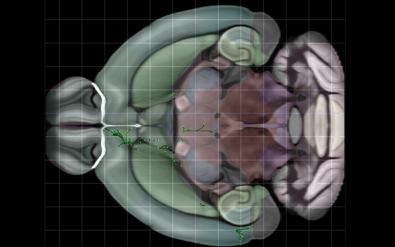

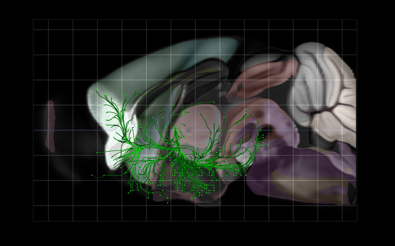

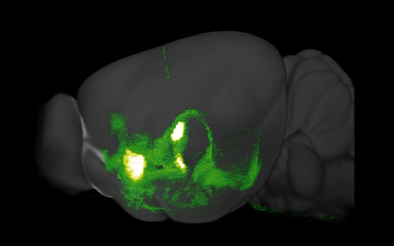

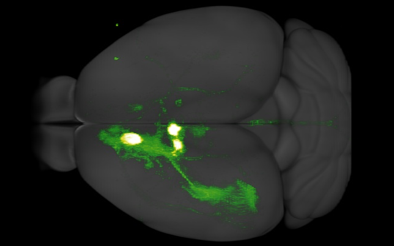

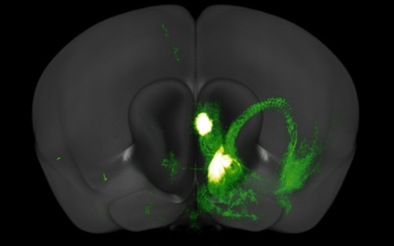

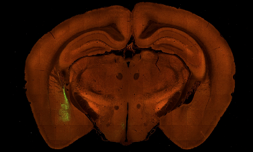

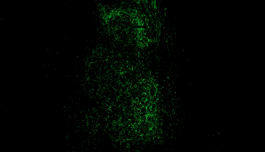

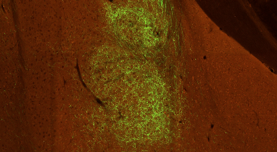

**B.**

**Figure S4. Three-dimensional reconstruction images of structural connections from Allen brain atlas database.**

(A) Virtual tractography between the cerebellum and the thalamus (B) Virtual tractography between the thalamus and the amygdala, and between the thalamus and the prelimbic cortex.
